# AKT1 mediates multiple phosphorylation events that functionally promote HSF1 activation

**DOI:** 10.1101/2020.08.31.275909

**Authors:** Wen-Cheng Lu, Ramsey Omari, Haimanti Ray, Imade Williams, Curteisha Jacobs, Natasha Hockaden, Matthew L. Bochman, Richard L. Carpenter

## Abstract

The heat stress response activates the transcription factor heat shock factor 1 (HSF1), which subsequently upregulates heat shock proteins to maintain the integrity of the proteome. HSF1 activation requires nuclear localization, trimerization, DNA binding, phosphorylation, and gene transactivation. Phosphorylation at S326 is an important regulator of HSF1 transcriptional activity. Phosphorylation at S326 is mediated by AKT1, mTOR, p38, and MEK1. Here, we observe that AKT1 activates HSF1 independent of mTOR. AKT2 also phosphorylated S326 of HSF1 but showed weak ability to activate HSF1. Similarly, mTOR, p38, and MEK1 all phosphorylated S326 but AKT1 was the more potent activator. Mass spectrometry showed that AKT1 also phosphorylated HSF1 at T142, S230, and T527 in addition to S326 whereas the other kinases did not. Subsequent investigation revealed that phosphorylation at T142 is necessary for HSF1 trimerization and that S230, S326, and T527 are required for HSF1 gene transactivation and recruitment of TFIIB and CDK9. This study suggests that HSF1 hyperphosphorylation is targeted and these specific residues have direct function in regulating HSF1 transcriptional activity.

## Introduction

Attenuation of cellular stressors is necessary for cell viability in unicellular and multicellular organisms. The discovery of a cellular response to heat stress was first observed in 1962 (1). Over two decades later, it was discovered that the protein heat shock factor 1 (HSF1) is the master regulator of the heat shock response (2, 3). Prior to heat stress-induced activation, HSF1 is sequestered in the cytoplasm within an inhibitory protein complex comprised of several molecular chaperone proteins (4–7). With the onset of heat stress, this complex dissociates and the HSF1 protein undergoes nuclear localization, trimerization, and phosphorylation leading to DNA binding at heat shock elements (HSEs) and gene transactivation (8–13). HSF1 target genes in response to heat stress include a large panel of chaperone proteins that safeguard the proteome from heat-induced denaturation (3, 14).

Dysregulation of HSF1 is associated with several disease phenotypes. Elevated HSF1 activity in multiple cancer types is a frequent occurrence (14–18). Loss of HSF1 delays or prevents formation of multiple cancer types (19–22). HSF1 appears to promote cancer by regulating genes distinct from its heat stress-induced gene targets (14) but has multiple functions to support tumor growth and progression (23). Conversely, loss of HSF1 is associated with neurodegenerative diseases, such as Huntington’s and Alzheimer’s diseases among others (24–31). HSF1 activity is impaired with aging but is critical for neuronal protein folding, neurotransmitter release, synapse formation, energy metabolism, and neuronal survival (32).

The majority of the functions of HSF1, both physiological and pathological, require a transcriptionally active HSF1. Transcriptionally active HSF1 exists in a trimer complex in the nucleus that is capable of binding to HSEs (8–13). HSF1 phosphorylation appears to strongly regulate the transcriptional activity of HSF1, particularly at S326 (8). Phosphorylation at S326 is routinely used as a marker of active HSF1 (8, 15, 23, 33–39). Since the initial observation that S326 phosphorylation affects heat stress-induced HSF1 activity (8), four kinases have been observed to phosphorylate S326 including mTORC1 (33, 37), MEK1 (38), p38 (34), and AKT1, which our work previously identified can phosphorylate S326 in breast cancer (15). However, the functional mechanism for how S326 phosphorylation promotes HSF1 transcriptional activity is not well understood. Furthermore, the relative contribution to HSF1 activity by these kinases is unknown. Additionally, it is not known if AKT1 contributes to heat stress-induced HSF1 activity or whether other AKT family members can activate HSF1. Our results indicate that AKT1 activates HSF1 most potently due to the unique ability of AKT1 to phosphorylate three additional residues beyond S326 that all contribute to different steps in the process of HSF1-driven transcription, including trimerization and gene transactivation. Interestingly, mTOR, p38, and MEK1 did not phosphorylate these additional sites.

## Methods and Materials

### Cell culture and reagents

All human breast cancer cell lines and human embryonic kidney cells (HEK293) used in this study were obtained from ATCC (Manassas, VA, USA), and maintained according to ATCC’s instructions. All chemicals were purchased from Sigma (St Louis, MO, USA) unless otherwise stated.

MK-2206 and rapamycin were purchased from Selleck Chemicals (Houston, TX). siAKT1, siAKT2, and non-target control (NT) oligonucleotide were purchased from Bioneer (Oakland, CA). All transfections were performed with cells in exponential growth using either X-tremeGENE siRNA (for siRNA) and X-tremeGENE HP DNA (for plasmids) (Roche, Indianapolis, IN, USA).

### Plasmids, transfection and luciferase assay

The FLAG-HSF-1 plasmid was purchased from Addgene (ID 32537, RRID:Addgene_32537), which was originally established by Dr Stuart Calderwood (40). The pcDNA3 Flag HA plasmid was purchased from Addgene (ID 10792, RRID:Addgene_10792), which was originally established by Dr William Sellers. The pcDNA3 Myr HA AKT1 plasmid was purchased from Addgene (ID 9008, RRID:Addgene_9008), which was originally established by Dr William Sellers (41). The pcDNA3 Myr HA AKT2 plasmid was purchased from Addgene (ID 9016, RRID:Addgene_9016), which was originally established by Dr William Sellers. The pcDNA3 Myr HA AKT3 plasmid was purchased from Addgene (ID 9017, RRID:Addgene_9017), which was originally established by Dr William Sellers. The pZW6 (CK2alpha) plasmid was purchased from Addgene (ID 27086, RRID:Addgene_27086), which was originally established by Dr David Litchfield (42). The pcDNA 3-CDK9 HA plasmid was purchased from Addgene (ID 14640, RRID:Addgene_14640), which was originally established by Dr Matija Peterlin (43). The pEGFP-C1-TFIIB plasmid was purchased from Addgene (ID 26675, RRID:Addgene_26675), which was originally established by Dr Sui Huang (44).

The heat shock element (HSE) firefly luciferase reporter was purchased from Promega (Madison, WI) and has two copies of the HSE promoter driving the luc2P gene. A Renilla luciferase expression vector, pRL-SV40P, was used to control for transfection efficiency. The pRL-SV40P was purchased from Addgene (ID 27163, RRID:Addgene_27163), which was originally established by Dr. Ron Prywes (45). 48hr after transfection, the cells were lysed and luciferase activity was measured using the Firefly and Renilla Luciferase Assay Kit (Promega). Relative promoter activity was computed by normalizing the Firefly luciferase activity against that of the Renilla luciferase.

### Immunoblotting and Immunoprecipitation (IP)

This was performed as we described previously (15, 46). Briefly, protein was isolated using RIPA buffer (50 mM Tris, 150 mM NaCl, 1 mM EDTA, 1% NP-40, 1% sodium deoxycholate, 1% SDS) for immunoblotting or with IP lysis buffer (20 mM Tris, 150 mM NaCl, 1 mM EDTA, 1 mM EGTA, 1 mM sodium pyrophosphate, 1 mM β-glycerophosphate, 1% Triton X-100) for lysate to be used for IP. Lysates were subjected to SDS-PAGE and subjected to immunoblotting with indicated antibodies. Antibodies used for immunoblotting included p-HSF1 (S326) (Abcam #76076, RRID:AB_1310328), HSF1 (CST #4356, RRID:AB_2120258), AKT1 (CST #2938, RRID:AB_915788), AKT2 (CST #3063, RRID:AB_2225186), AKT3 (CST #3788, RRID:AB_2242534), beta-actin (BD Biosciences #612656, RRID:AB_2289199), GAPDH (CST #2118, RRID:AB_561053), GST (CST #2625, RRID:AB_490796), Flag (Sigma #3165, RRID:AB_259529), CDK9 (CST #2316, RRID:AB_2291505), and GFP (CST #2956, RRID:AB_1196615). Antibodies used for IP were FLAG resin beads (Sigma #A2220) and control mouse IgG beads (Sigma #A0919).

### Kinase Assay and Mass Spectrometry

Recombinant AKT1, recombinant AKT2, recombinant HSF-1, recombinant p38, recombinant mTOR, recombinant MEK and recombinant ERK were purchased from Sigma. Indicated amounts of recombinant GST-HSF-1 (0.1–0.4 mg) was incubated with or without 0.1 mg recombinant kinases for up to 120 min at 30°C in the presence of ATP in kinase assay buffer (100 mM HEPES, 50 mM MgCl_2_, 50 mM MnCl_2_). Samples were then boiled and subjected to SDS–PAGE and WB with indicated antibodies or subjected to mass spectrometry for detection of post-translational modifications.

Mass spectrometry for post-translational modifications was performed at the Laboratory for Biological Mass Spectrometry at Indiana University (Bloomington, IN). Protein samples were dissolved in 8M urea, 100mM ammonia bicarbonate (pH8.0) solution. Samples were incubated for 1 hr at 56°C with 10 mM TCEP to reduce cysteine residue side chains. The alkylation of cysteine residue side chains was proceeded for 45 min at room temperature in the dark with 20 mM iodoacetamide. For chymotrypsin digestion, 1 μg of chymotrypsin was added to the diluted sample and the samples were digested at 25°C overnight. Desalted peptides were injected into an Easy-nLC 100 HPLC system coupled to an Orbitrap Fusion Lumos mass spectrometer (Thermo Scientific, Bremen, Germany). Peptide samples were loaded onto an Acclaim PepMapTM 100 C18 trap column (75 μm × 20 mm, 3 μm, 100 Å) in 0.1% formic acid. The peptides were separated using an Acclaim PepMapTM RSLC C18 analytical column (75 μm × 150 mm, 2 μm, 100 Å) using an acetonitrile-based gradient (Solvent A: 0% acetonitrile, 0.1% formic acid; Solvent B: 80% acetonitrile, 0.1% formic acid) at a flow rate of 300 nl/min. A 30 min gradient was as follows: 0-0.5 min, 2-8% B; 0.5-24 min, 7-38% B; 24-26 min, 40-100% B; 26-30 min, 100% B, followed by re-equilibration to 2% B. The electrospray ionization was carried out with a nanoESI source at a 260 °C capillary temperature and 1.8 kV spray voltage. The mass spectrometer was operated in data-dependent acquisition mode with mass range 350 to 2000 m/z. The precursor ions were selected for tandem mass (MS/MS) analysis in Orbitrap with 3 sec cycle time using HCD at 30% collision energy. Intensity threshold was set at 2.5e4. The dynamic exclusion was set with a repeat count of 1 and exclusion duration of 30 s. The resulting data was searched in Protein Prospector (http://prospector.ucsf.edu/prospector/mshome.htm) against Human HSF1 sequence (Q00613). Carbamidomethylation of cysteine residues was set as a fixed modification. Protein N-terminal acetylation, oxidation of methionine, protein N-terminal methionine loss, pyroglutamine formation, phosphorylation on STY were set as variable modifications. A total of two variable modifications were allowed. Chymotrypsin digestion specificity with two missed cleavage was allowed. The mass tolerance for precursor and fragment ions was set to 5 ppm and 10 ppm respectively. Peptide and protein identification cut-off scores were set to 15 and 22, respectively.

### HSF1 Trimerization and Gel Filtration Chromatograhy

Cells were transfected with HSF1-Flag for 48 hrs. Cells were then fixed by adding 1 mM ethylene glycol bis-succinimidylsuccinate (Thermo Scientific, Waltham, MA) for 30 min at room temperature (RT) for crosslinking. The crosslinking reaction was quenched by addition of 10 mM Tris, pH 7.5, for 15 min at RT. Protein was isolated using RIPA buffer and the resulting lysate underwent SDS-PAGE and subjected immunoblotting with antibodies to HSF1.

Gel filtration chromatography was performed using fixed cells as described above for trimerization. Total protein was isolated using RIPA and the resulting lysate was loaded onto a Superdex 200 Increase 10/300 GL column (Cytivia, Marlborough, MA) that was pre-equilibrated in running buffer (50 mM Tris, pH 8.0, 150 mM NaCl, and 1% NP-40). Proteins were eluted with 2 column volumes of running buffer and captured as 2-mL fractions. All chromatography steps were performed at 4°C, and the column was calibrated using running buffer and a Gel Filtration Markers Kit (Sigma). Samples (10 μL) of each fraction that eluted within the molecular standard range of 29-669 kDa were run on 4-15% Mini-PROTEAN TGX (Bio-Rad, Hercules, CA) SDS-PAGE gels at 15 V/cm. The proteins were transferred to nitrocellulose membranes at 4°C and blocked with non-fat milk in TBST at room temperature. The blots were probed with 1:1000 anti-HSF1 antibody and visualized with IRDye 800CW Goat anti-Rabbit secondary antibody (LI-COR, Lincoln, NE) and a LI-COR Odyssey imager.

### Chromatin immunoprecipitation (ChIP)

This was performed using a ChIP Assay Kit (Millipore-Upstate, Billerica, MA, USA) as we described previously (15). Rabbit polyclonal HSF1 antibody was used for IP (CST #4356). Primer sequences for ChIP-qPCR are listed in **Table 1**.

**Table 1:**
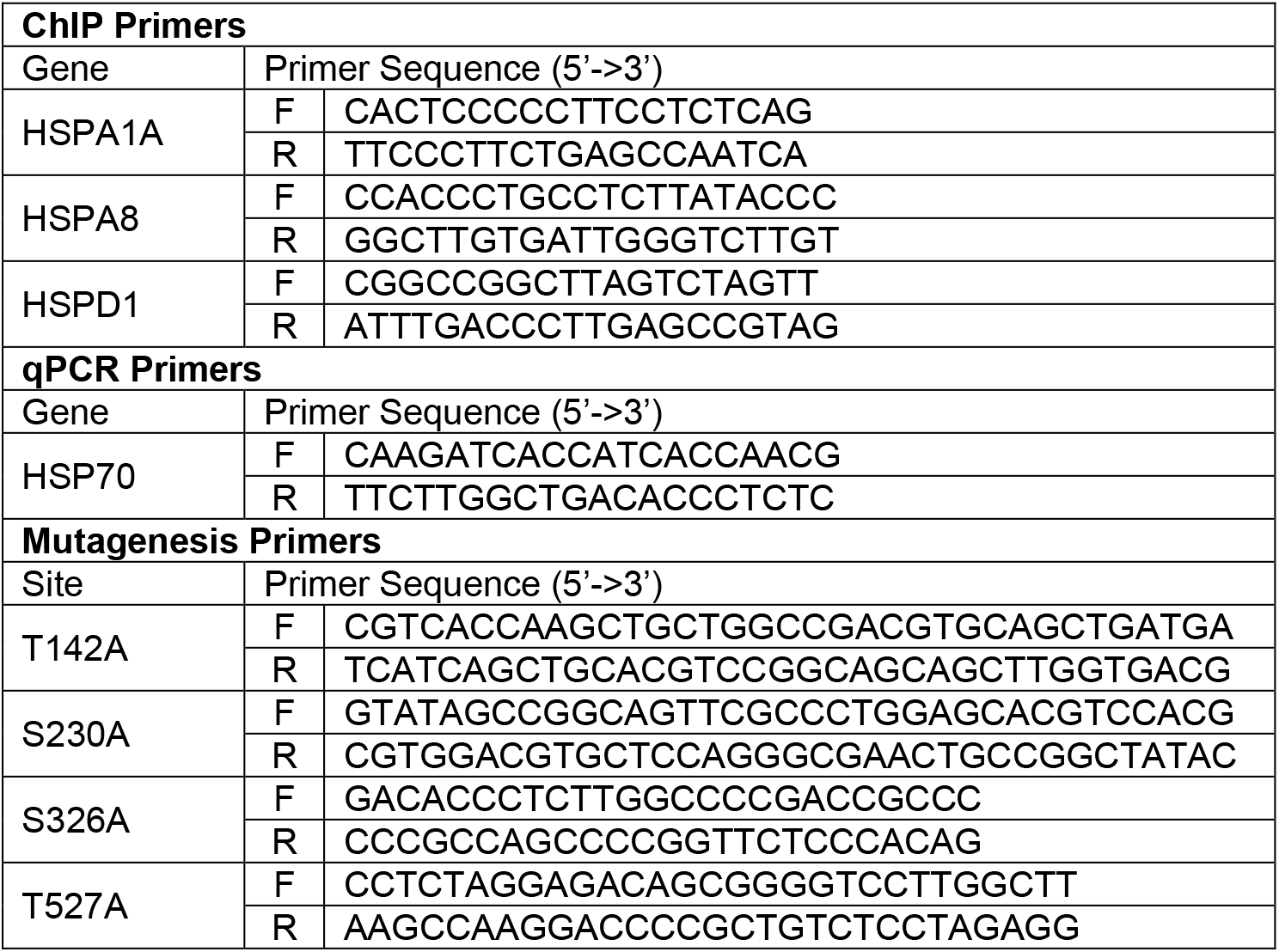
Primers.

### qRT-PCR analysis

For analysis of mRNA levels, total RNA was collected from cells using an RNA isolation kit (Zymo, Santa Cruz, CA). Total RNA underwent reverse transcription (RT) using an RT kit (Applied Biosystems, Foster City, CA) that uses a mixture of oligodT and random primers. The resulting cDNA underwent qPCR using gene-specific primers with the SYBR Green qPCR Assay system (Applied Biosystems). Primer sequences for qPCR are listed in **Table 1**. Relative mRNA levels were computed using the ΔΔCt method.

### Gene Set Enrichment Analysis (GSEA)

GSEA was done as previously described (46, 47). Gene Cluster Text file (.gct) was generated from the TCGA BRCA or TCGA COADREAD datasets. Categorical class file (.cls) was generated based on a signature for HSF1 activity (14). The Gene Matrix file (.gmx) was generated using published gene signatures for AKT1 (48), AKT2 (48), p38 (47), mTORC1 (47), and MEK (49). The number of permutations was set to 1000 and the chip platform for TCGA gene lists was used. Normalized enrichment scores (NES) and adjusted p-values are reported.

### Mutagenesis

Generation of mutant HSF1-T142A, HSF1-S230A, HSF1-S326A and HSF1-T527A were carried out using a QuikChange II XL Site-Directed Mutagenesis kit (Agilent Technologies, Santa Clara, CA) as per the manufacturer’s instructions. Primers used for mutagenesis are listed in **Table 1**. Mutation was confirmed by sequencing.

### Statistical analysis

Data are presented as mean ± SEM. Unpaired *t*-tests and ANOVA with Tukey’s post-hoc test were used for statistical analysis where appropriate. A *p* value of <0.05 was considered statistically significant.

## Results

### AKT1 and AKT2 phosphorylate HSF1 at S326 but only AKT1 activates HSF1

We previously identified AKT1 as an upstream regulator of HSF1 activity via phosphorylation of HSF1 at S326 (15). AKT1 has well-known oncogenic functions (50, 51). AKT2 and AKT3 also have pro-tumor functions (52–55) but whether HSF1 can be regulated by these isoforms is unknown. To assess this possibility, either AKT1, AKT2, or AKT3 were introduced into multiple cell lines along with a luciferase reporter driven by the presence of two heat shock elements (HSEs). Consistent with our previous findings, AKT1 strongly induced HSF1 activity (**Fig 1A-1C**). AKT2 promoted a significant increase in HSF1 activity but the effect was modest while AKT3 had no significant effect on HSF1 activity (**Fig 1A-1C**). To determine the ability of the AKT isoforms to phosphorylate HSF1 at S326, a key phosphorylation event critical for HSF1 activity (8), an in vitro kinase assay followed by immunoblotting was performed with each AKT isoform. Both AKT1 and AKT2 phosphorylated S326 of HSF1 at similar levels but AKT3 showed no capability for phosphorylating S326 (**Fig 1D**). These results seem contradictory because AKT2 phosphorylated S326 but the level of phosphorylation far exceeded the HSF1 activity induced by AKT2 (**Fig 1A-1D**). However, we also observed that AKT2 did not phosphorylate S326 as strongly in cells (**Fig 1E**), suggesting AKT2 has the capability to phosphorylate HSF1 but this is more limited within living cells. AKT1 is also a well-known activator of mTOR (56, 57) and mTOR has been shown to phosphorylate HSF1 at S326 in response to proteotoxic stress (33, 37). Therefore, to determine whether AKT1-driven HSF1 activity and S326 phosphorylation is mediated by mTOR, cells were treated with increasing concentrations of rapamycin and subjected to the HSE luciferase reporter assay or immunoblotting with p-HSF1 S326 antibodies. Results indicated that rapamycin had no effect on HSF1 activity (**Fig 1F**) or on S326 phosphorylation (**Fig 1G**) driven by AKT1. We further assessed whether the AKT isoforms affected downstream activation steps of HSF1 including trimerization and DNA binding. Introduction of AKT1 was observed to promote a significant increase in trimerization, although the AKT1-driven trimerization was not as much as heat stress (**Fig 2A-B**). AKT2 and AKT3 promoted on minimal amounts of HSF1 trimerization (**Fig 2A-B**). Similarly, introduction of AKT1 promoted a significant increase of HSF1 binding to putative targets HSPA1A (HSP70), HSPA8 (HSP73), and HSPD1 (HSP60) whereas AKT2 and AKT3 had no significant effect (**Fig 2C**). We did not observe any effect of AKT1 on HSF1 protein half-life (**Suppl. Fig. 1**). These results indicate AKT1 is the primary AKT isoform capable of activating HSF1 with AKT2 having modest effects in cells.

**Figure 1.**
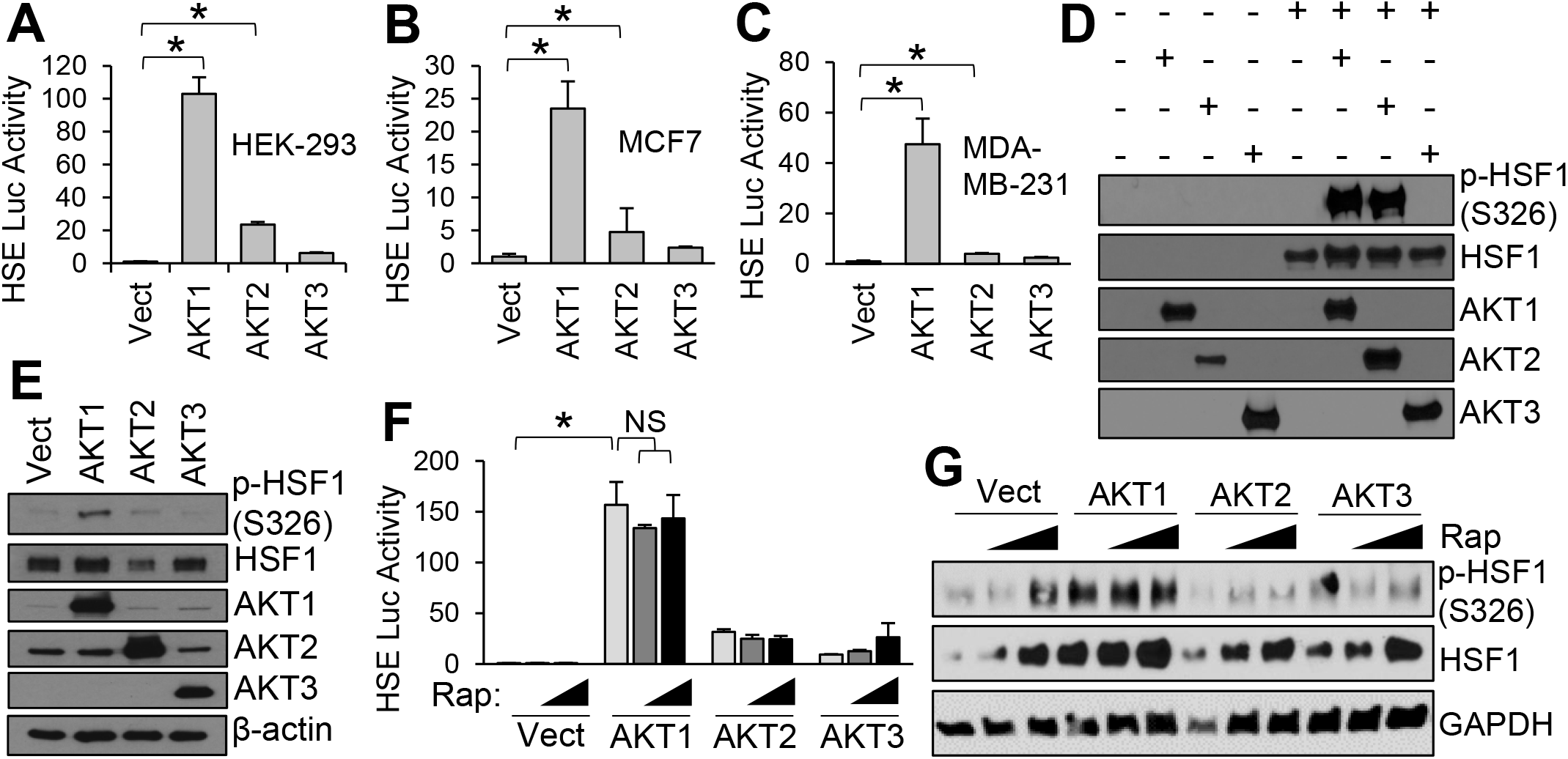
AKT1 Phosphorylates HSF1 on S326 and induces HSF1 activity independent of mTORC1. A-C) Indicated cell lines were transfected with vector, AKT1, AKT2, or AKT3 along with the HSE luciferase reporter for 48 hrs. Firefly/Renilla ratios were compared across groups. D) Recombinant proteins were incubated at 30°C for 2 hrs in the presence of ATP and subsequently subjected to SDS-PAGE and immunoblotting with indicated antibodies. E) HEK-293 cells were transfected with vector, AKT1, AKT2, or AKT3 for 48 hrs. Cell lysates were subjected to SDS-PAGE and immunoblotting with indicated antibodies. F-G) HEK-293 cells were transfected with vector, AKT1, AKT2, or AKT3 in the presence or absence of rapamycin (10-20 nM). Cell lysates were subjected to firefly reporter assay (F) or immunblotting with indicated antibodies (G). Experiments were completed in triplicate and analyzed using one-way ANOVA and Tukey’s post-hoc test. Data is presented as mean ± SEM. *P<0.05.

**Figure 2.**
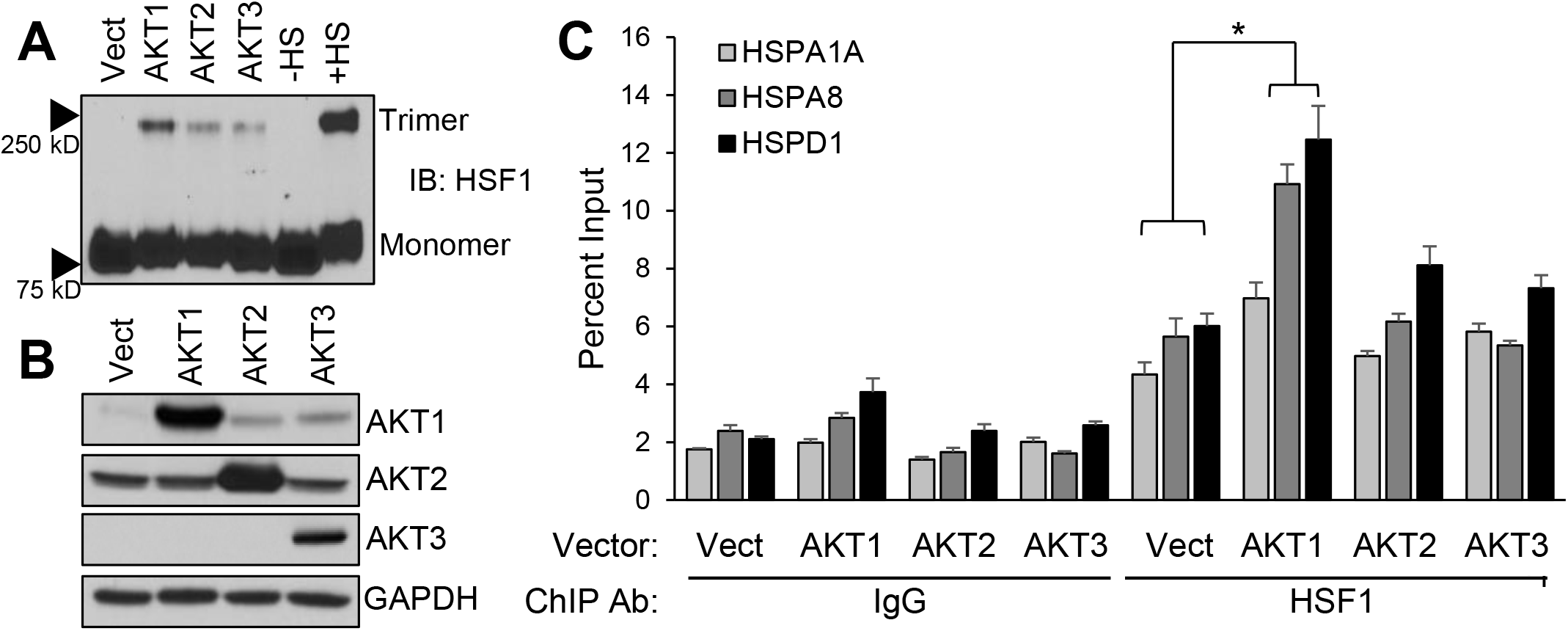
AKT1 promotes HSF1 trimerization and DNA binding. A-B) HEK-293 cells were transfected with vector, AKT1, AKT2, or AKT3 for 48 hrs. Cells were fixed with 1 mM EGS for 30 min and lysates were subjected to SDS-PAGE and immunoblotting for HSF1 trimers (A) and AKT isoforms (B). C) HEK-293 cells were transfected with vector, AKT1, AKT2, or AKT3 for 48 hrs. Cells were fixed with formaldehyde, lysed, and sonicated prior to ChIP with either control IgG or HSF1 antibodies. ChIP product was subjected to qPCR for indicated gene promoters. Experiments were completed in triplicate and analyzed using one-way ANOVA and Tukey’s post-hoc test. Data is presented as mean ± SEM. *P<0.05.

### AKT1 is required for full activation of HSF1 in response to heat stress

Activation of the PI3K-AKT pathway in response to heat stress has previously been observed as far back as 2000 (58, 59). However, the precise role of this pathway in heat stress-induced activation of HSF1 has not been fully explored. Our previous work indicated that HSF1 can be activated by AKT1 via phosphorylation at S326 in breast cancer cells (15). To assess whether AKT signaling is crucial for heat-induced activity of HSF1, cells were subjected to heat stress in the presence or absence of the pan-AKT inhibitor MK-2206. Results indicated that blocking AKT activity suppressed HSF1 activity in response to heat stress (**Fig 3A**). Heat stress was also observed to induce HSF1 phosphorylation and knockdown of AKT1 significantly reduced HSF1 phosphorylation in response to heat (**Fig 3B**). Loss of AKT2 did not significantly affect HSF1 phosphorylation in response to heat stress (**Fig 3B**). Similarly, loss of AKT1 reduced HSF1 reporter activity by more than 50% in response to heat stress (**Fig 3C**). Interestingly, loss of AKT2 reduced HSF1 activity by a small percentage (∼20%) but this was not statistically significant (**Fig 3C**). In accordance with the phosphorylation results, loss of AKT1 significantly reduced levels of HSP70, a direct HSF1 gene target, in response to heat but loss of AKT2 had no significant effect (**Fig 3D**). While previous investigations primarily focused on PI-3K inhibitors to determine the relevance of this pathway to heat stress responses (59), these data indicate that AKT1 is an important factor for the activation of HSF1 in response to heat stress.

**Figure 3.**
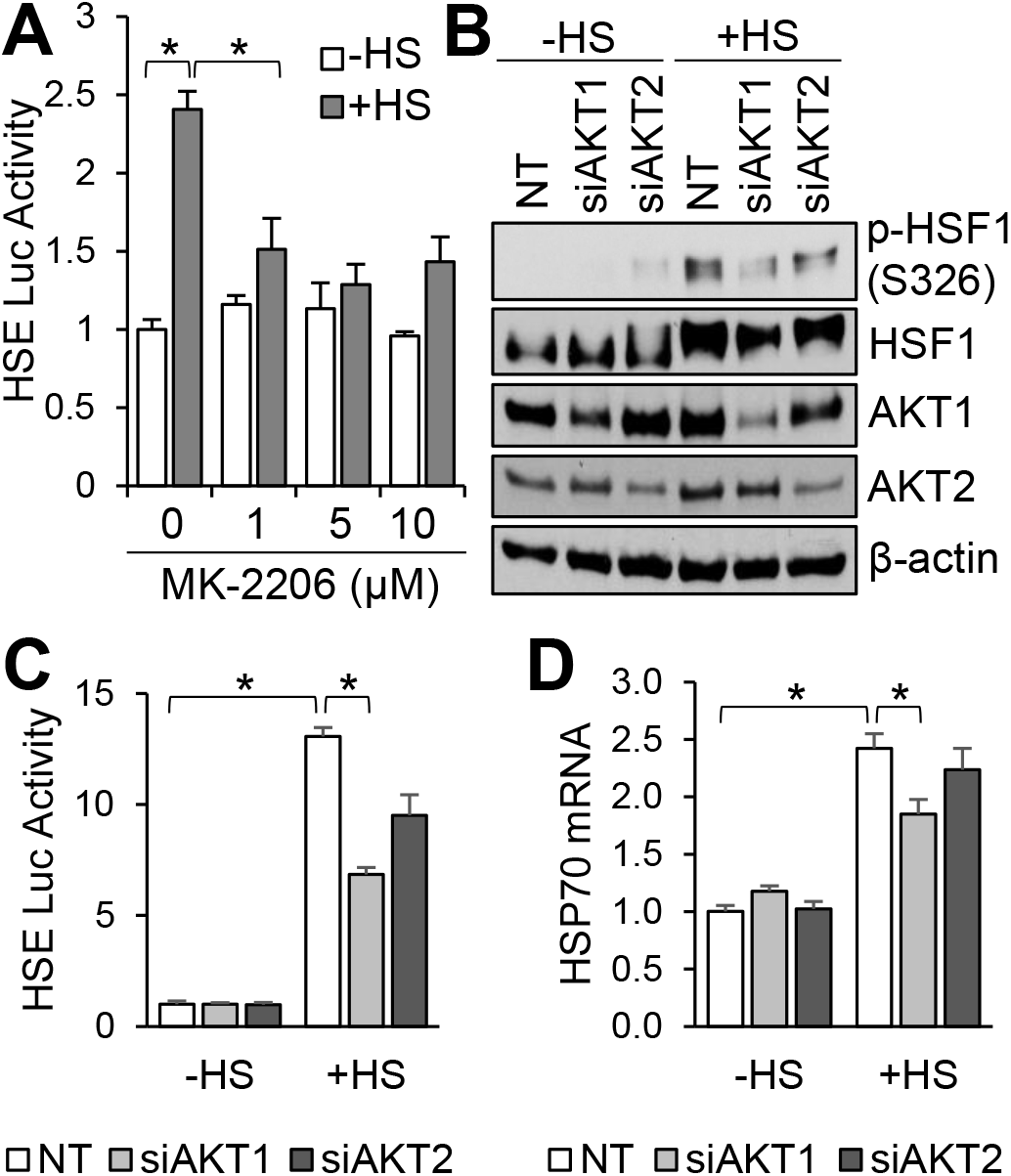
AKT1 is required for full HSF1 activation in response to heat stress. A) HEK-293 cells were transfected with the HSE luciferase reporter for 48 hrs and cells were then pre-treated with MK-2206 at the indicated concentrations for 1 hr and cells were then incubated at 42°C for 1 hr. Firefly/Renilla ratios were compared across groups. B-D) HEK-293 cells were transfected with non-targeting (NT), AKT1, or AKT2 siRNA for 48 hrs followed by incubation at 42°C for 1 hr. Cells were then collected and subjected to immunoblotting (B), luciferase reporter assay (C), or qPCR (D). Experiments were completed in triplicate and analyzed using one-way ANOVA and Tukey’s post-hoc test. Data is presented as mean ± SEM. *P<0.05.

### AKT1 is a more potent activator of HSF1 than other known activator kinases

Our above results indicated that both AKT1 and AKT2 can phosphorylate HSF1 at S326 but AKT1 is capable of promoting a substantially higher transcriptional activity of HSF1 (**Fig 1–2**). To test the importance of S326 phosphorylation, the three other kinases mTORC1 (33, 37), MEK1 (38), and p38 (34) that have previously been shown to phosphorylate HSF1 at S326 were also tested using an in vitro kinase assay. Results were similar to that which was previously shown in that all of these kinases phosphorylated HSF1 at S326 with AKT1 and mTOR showing the highest phosphorylation (**Fig 4A**). However, due to this being in vitro assay, it is unclear whether this pattern is reflective of the relative phosphorylation of S326 by these kinases in cells. Therefore, all of these kinases were introduced into cells and all kinases increased S326 phosphorylation of HSF1 to similar levels suggesting in cells they all have a similar phosphorylation capability (**Fig 4B**). Interestingly, however, AKT1 was more potent than these kinases in activating HSF1 transcriptional activity (**Fig 4C**). Similarly, knockdown of AKT1 significantly reduced heat stress-induced HSF1 activity along with knockdown of mTOR whereas knockdown of MEK1 and p38δ had little effect on heat stress-induced HSF1 activity (**Fig. 4D**).

**Figure 4.**
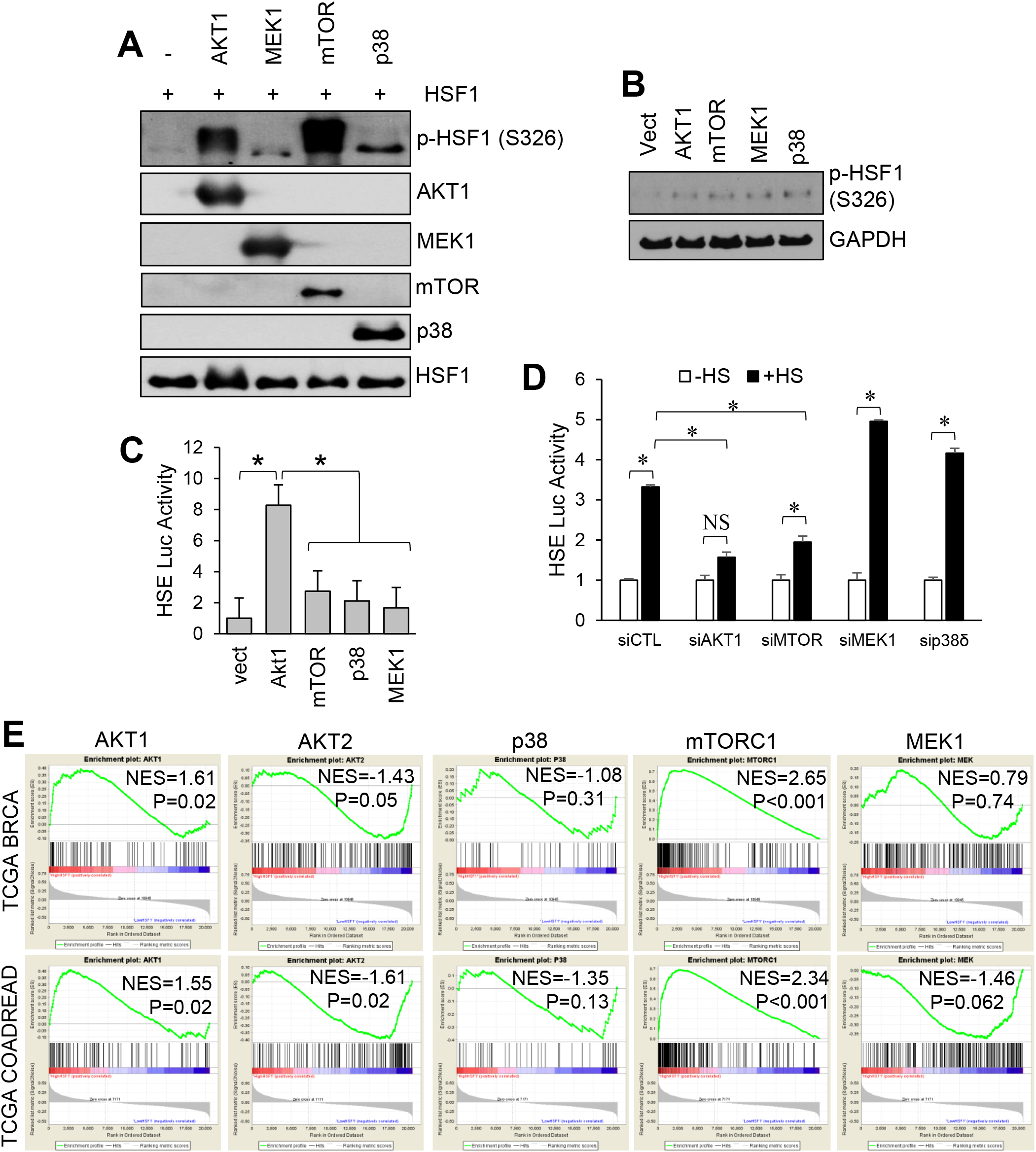
AKT1 induced HSF1 activity stronger than other known activating kinases despite similar induction of S326 phosphorylation. A) Recombinant proteins were incubated at 30°C for 2 hrs in the presence of ATP and subsequently subjected to SDS-PAGE and immunoblotting with indicated antibodies. B-C) HEK-293 cells were transfected with vector, AKT1, mTOR, MEK1, or p38 for 48 hrs. Cells were collected and subjected to immunoblotting with indicated antibodies (B) or reporter luciferase assay (C). D) HEK-293 cells were transfected with non-targeting control siRNA or siRNA directed to AKT1, mTOR, MEK1, or p38δ along with an HSE luciferase reporter. Cells were then incubated at 42°C for 1 hr and subsequently collected and subjected to the luciferase reporter assay. E) Gene set enrichment analysis (GSEA) was performed using data from TCGA for breast (upper row) and colorectal cancers (lower row). Patients from these datasets were separated into high and low HSF1 activity groups using previously published gene signature for HSF1 activity. GSEA was then performed on these two groups using activity signatures for AKT1, AKT2, p38, mTORC1, or MEK1. Normalized enrichment score (NES) is reported and calculated GSEA p-value. Experiments were completed in triplicate and analyzed using one-way ANOVA and Tukey’s post-hoc test. Data is presented as mean ± SEM. *P<0.05. Data in D has normalized enrichment score (NES) reported and calculated GSEA p-value.

To further assess the correlation between these kinases and HSF1 activity, gene set enrichment analysis (GSEA) was performed (47, 60). This analysis was performed using TCGA datasets for breast cancer (BRCA) and colorectal adenocarcinoma (COADREAD) wherein patients for each dataset were scored for HSF1 activity using the HSF1 CaSig (14). The patients in each dataset were then separated into high HSF1 activity (upper 50%) or low HSF1 activity (lower 50%) based on HSF1 signature score and these two groups were set as the two phenotypes for GSEA. Enrichment in these two groups of HSF1 activity were performed using signatures for activity of AKT1 (61, 62), AKT2 (48), p38 (47), mTORC1 (63), and MEK1 (49). Results indicated that signatures for AKT1 and mTORC1 were significantly enriched in high HSF1 activity tumors (**Fig. 4E**), suggesting a strong correlation between the activity of these kinases and HSF1 activity. The signature for AKT2 was significantly enriched in the low HSF1 activity group in the COADREAD patients and borderline significant in the BRCA patients, suggesting a possible weak negative correlation between AKT2 and HSF1 activities. There was no significant enrichment toward either high or low HSF1 activity for p38 or MEK1 signatures. Taken together, these data indicate a lack of consistency between HSF1 phosphorylation at S326 by this group of kinases and the resulting HSF1 transcriptional activity.

### AKT1 phosphorylates additional residues aside from S326 that are critical for HSF1 activity

Data from Figure 4 indicates that several kinases can phosphorylate HSF1 at S326, its canonical activating phosphorylation event (8). However, phosphorylation by these different kinases resulted in varying degrees of HSF1 transcriptional activity with AKT1 promoting higher activity than the other kinases. To resolve this apparent contradiction, we reasoned that AKT1 may be phosphorylating additional residues that support HSF1 activation. To test this hypothesis, recombinant HSF1 was subjected to a kinase assay with either AKT1, AKT2, p38, mTOR, or MEK1, as performed in Figures 1D and 4A, followed by mass spectrometry to identify novel phosphorylation sites (**Fig 5A**). Results from this phosphorylation scan are summarized in **Fig 5B** and **Supplemental Table 1**, which illustrates the phosphorylation sites outside of the known target of S326. AKT1 phosphorylated T142, S230, and T527. Interestingly, these sites were entirely unique to AKT1 as only MEK1 was seen to phosphorylate any of these sites by phosphorylating T527. All four of these sites, including S326, are conserved across several mammalian species (**Fig 5C**). Phosphorylation at T142 has only been reported once and this was mediated by CK2 (64). Phosphorylation at S230 has been previously reported and was shown to promote inducible HSF1 activity (65). Phosphorylation at T527 has never been reported. Phospho-null mutations to alanine at each of these sites individually resulted in some variable effects with S326A and S230A having no significant effect of HSF1 activity in response to heat stress (**Fig. 5D**). T142A and T527A individually showed a small but significant decrease in HSF1 activity in response to heat stress (**Fig 5D**). However, when all four of these sites were mutated to alanine, HSF1 no longer responded to heat stress (**Fig 5E**). Phospho-mimic mutants for these residues showed highly variable results with little evidence that T/S->E mutations were sufficiently mimicking phosphorylation, thus rendering these mutants unusable as a reliable phospho-mimic (**Suppl. Fig. 2**). These results indicate that AKT1 modifies HSF1 in a unique pattern by phosphorylating T142, S230, and T527 in addition to S326. This unique pattern of phosphorylation may mechanistically explain why AKT1 promotes HSF1 activity more potently than these other kinases.

**Figure 5.**
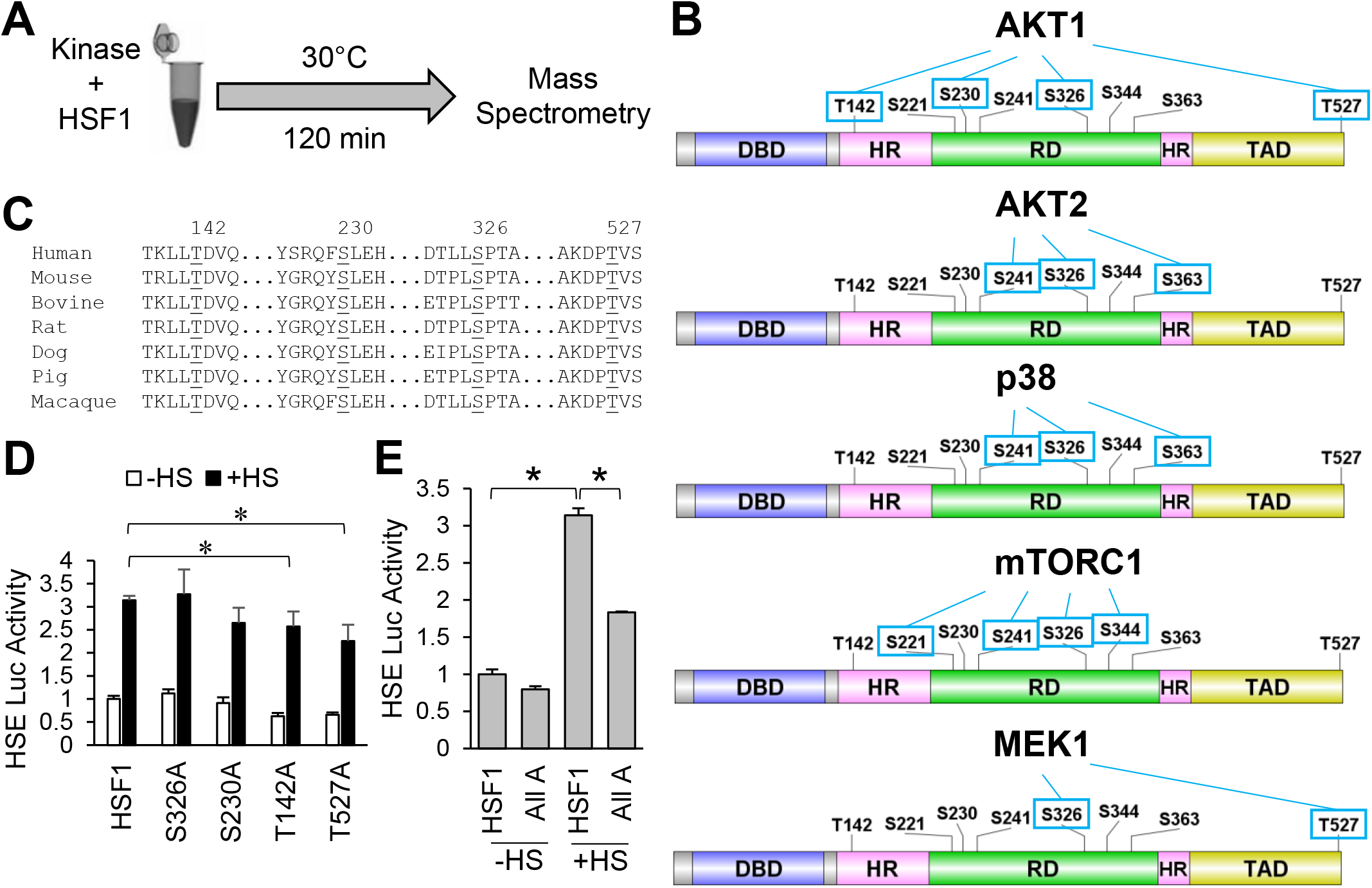
AKT1 phosphorylates four residues that regulate the HSF1 response to heat stress. A) Workflow for identification of kinase-specific phosphorylation sites. B) Pattern of HSF1 phosphorylation for each kinase tested by mass spectrometry. C) Conservation of four sites phosphorylated by AKT1 across mammalian species. D) HEK-293 cells were transfected with WT-HSF1 or single phospho-null HSF1 mutants for AKT1 phosphorylation sites along with the HSE reporter for 48 hrs followed by incubation at 42°C for 1 hr. Firefly/Renilla ratios were compared across groups. E) HEK-293 cells were transfected with WT-HSF1 or an HSF1 mutant with all four AKT1 phosphorylation sites with phospho-null mutations for 48 hrs followed by incubation at 42°C for 1 hr. Firefly/Renilla ratios were compared across groups. Experiments were completed in triplicate and analyzed using one-way ANOVA and Tukey’s post-hoc test. Data is presented as mean ± SEM. *P<0.05.

### Phosphorylation at T142 enhances trimerization of HSF1

AKT1 phosphorylates HSF1 at multiple residues in addition to S326 (**Fig 5**). Should these additional phosphorylation sites mediate the additional HSF1 activity observed in the presence of AKT1, it stands to reason that these sites functionally participate in the activation process of HSF1. Therefore, the role of each of these four phosphorylation sites in the process of HSF1 activation was investigated. Threonine 142 sits in the first leucine-zipper domain, also referred to as the heptad repeat domain (32), and this domain is responsible for trimerization of HSF1 (11, 13, 66)). Considering T142 is located in this domain, it was tested whether phosphorylation at this site affected trimerization. Introduction of AKT1 with wild type (WT)-HSF1 resulted in an increase in trimerization, which was lost in the presence of the T142A phospho-null mutant (**Fig 6A**). To test whether T142 regulation of trimerization was AKT1-specific, HSF1 trimerization was also tested in the presence of CK2, which has previously been observed to phosphorylate T142 (64). In the presence of CK2, WT-HSF1 also showed a significant increase in trimerization but T142A-HSF1 did not undergo trimerization in the presence of CK2 (**Fig 6B**). These results likely indicate that phosphorylation at T142 is an important regulatory step of HSF1 trimerization and is not AKT1-specific. To further assess the oligomeric state of HSF1 and the role of T142, gel filtration chromatography was performed on lysates expressing either WT-HSF1 or T142A-HSF1 with or without the presence of AKT1. When fractions from WT-HSF1-expressing cells were blotted for HSF1, it was found to elute in the lower molecular weight range, consistent with the molecular weight that HSF1 monomers migrate at approximately 70-80 kDa (**Fig. 6C**). In the presence of AKT1, HSF1 was found in earlier-eluting fractions with a predicted molecular weight range indicative of trimers. However, T142A-HSF1 in the presence of AKT1 eluted at lower molecular weights, indicative of monomers. These results together indicate that phosphorylation at T142 is necessary for effective trimerization of HSF1.

**Figure 6.**
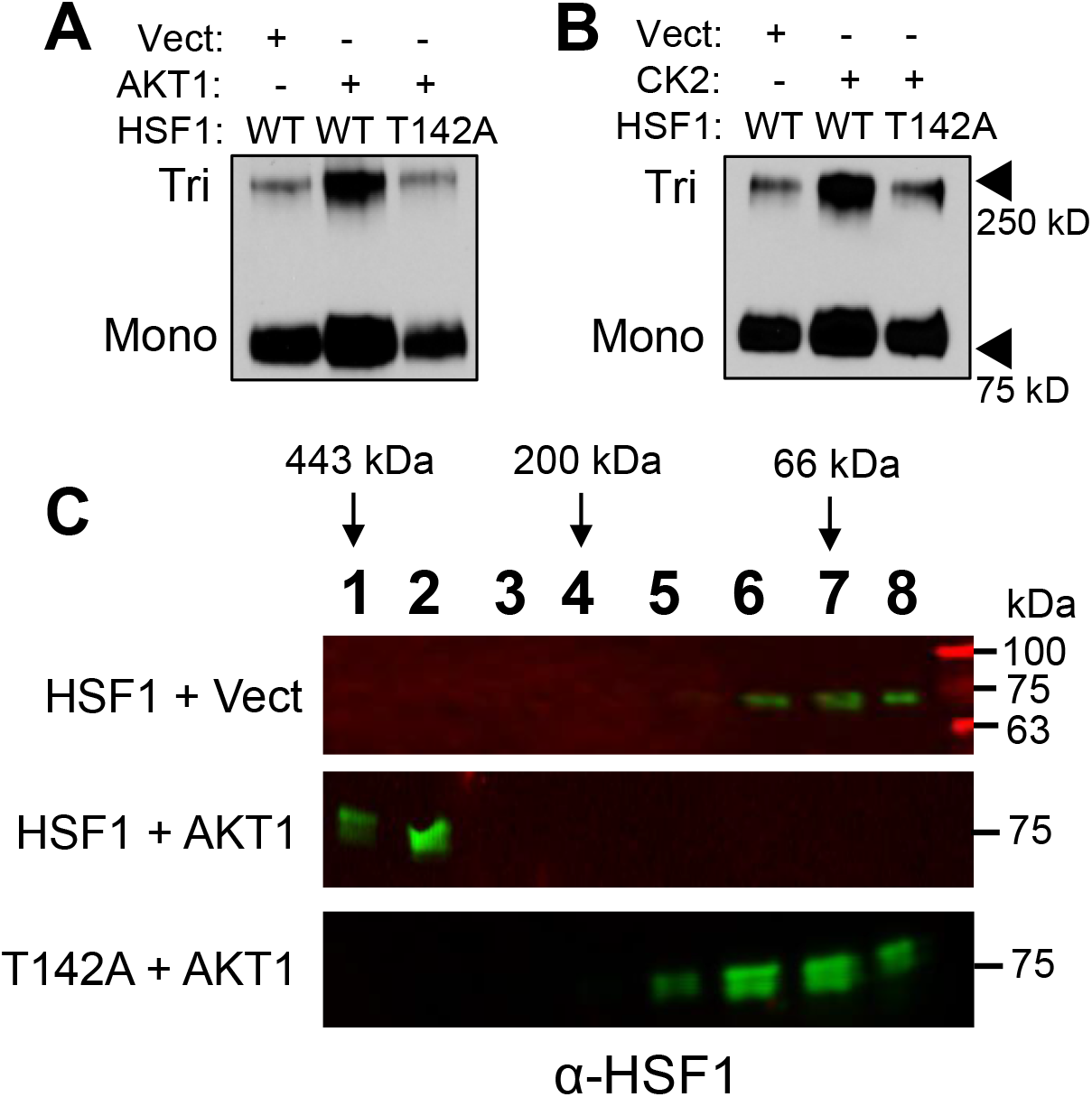
Phosphorylation at T142 enhances HSF1 trimerization. A-C) HEK-293 cells were transfected with WT-HSF1 or T142A-HSF1 in the presence or absence of AKT1 (A, C) or CK2 (B) for 48 hrs. Cells were fixed with 1 mM EGS for 30 min and lysates were subjected to SDS-PAGE and immunoblotting for HSF1 trimers. Samples in C were subjected to size exclusion chromatography and fractions underwent immunoblotting with HSF1 antibodies to identify HSF1 complex sizes according to the molecular weight of the extracted fractions. A and B were completed in triplicate and analyzed using one-way ANOVA and Tukey’s post-hoc test. Data is presented as mean ± SEM. *P<0.05.

### Phosphorylation at S230, S326, and T527 enhances HSF1 association with general transcriptional factors

The human HSF1 protein has 529 residues, of which 103 are residues potentially capable of phosphorylation by being tyrosine (n=8), serine (n=70), or threonine (n=25). A total of 39 of these residues have been previously shown to be phosphorylated either by non-biased screening using mass spectrometry or low-throughput methods (67). Our analysis identified T527 as a phosphorylated residue in the presence of AKT1 (**Fig 5**), which is a novel site never before seen phosphorylated. Considering this residue has never been observed as phosphorylated, the role of this phosphorylation event in HSF1 activation is unknown. Since this residue resides in the transactivation domain (TAD) of HSF1, it was hypothesized that it may contribute to the function of the TAD in recruiting transcriptional machinery for initiation of transcription. HSF1 has previously been shown to have physical association with CDK9 as part of the p-TEFb complex, TFIIB, and TATA binding protein (TBP) (68–72). To initially screen if phosphorylation at T527 affects HSF1 interaction with these proteins, either WT- or T527A-HSF1 were expressed in cells in the presence of CDK9, TFIIB, or TBP followed by HSF1 activity assessed with an HSE reporter. HSF1 activity was significantly reduced with T527A phospho-null mutant in the presence of CDK9 and TFIIB but not TBP (**Fig 7A-C**). We observed similar results with mRNA levels of HSP70, a direct transcriptional target of HSF1 (**Suppl. Fig. 3A-C**), suggesting this phosphorylation site may be involved in interaction with CDK9 and TFIIB. Co-immunoprecipitation revealed a loss of interaction between T527A-HSF1 with CDK9 and TFIIB (**Fig 7D-E**), suggesting phosphorylation at this site is critical for this protein complex formation.

**Figure 7.**
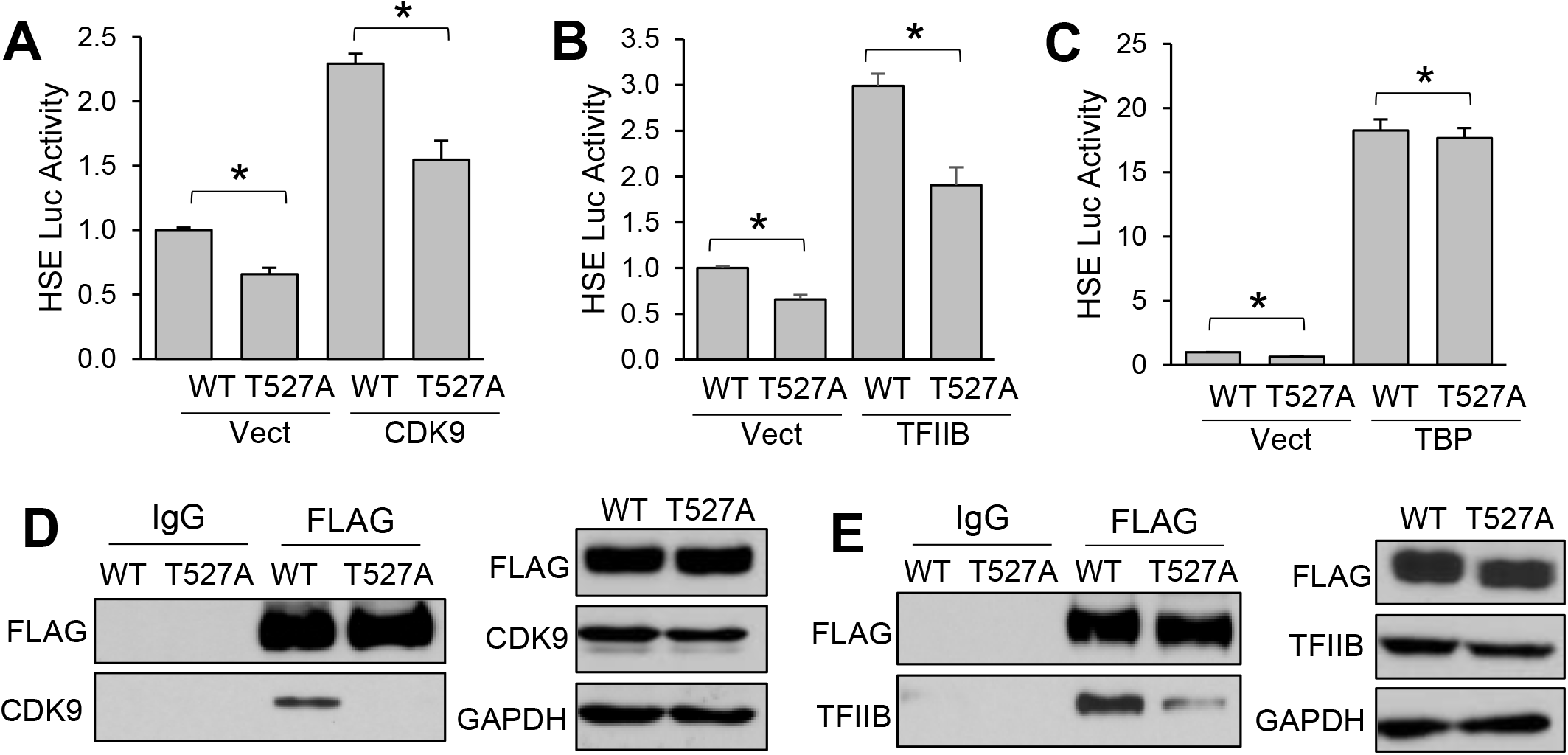
Phosphorylation at T527 enhances HSF1 association with general transcription factors. A-C) HEK-293 cells were transfected with WT-HSF1 or T527A-HSF1 in the presence or absence of CDK9 (A), TFIIB (B), or TBP (C) along with the HSE luciferase reporter for 48 hrs. Firefly/Renilla ratios were compared across groups. D-E) HEK-293 cells were transfected with FLAG-tagged WT-HSF1 or T527A-HSF1 for 48 hrs. Cell lysates were subjected to immunoprecipitation with control IgG or FLAG antibodies and subjected to immunoblotting with indicated antibodies. A-C were completed in triplicate and analyzed using one-way ANOVA and Tukey’s post-hoc test. Data is presented as mean ± SEM. *P<0.05.

Phosphorylation at S326 to upregulate HSF1 activity is well documented (8, 15, 33, 34, 37, 38). Phosphorylation at S230 has also been observed to promote HSF1 transcriptional activity (65). However, the precise function of these phosphorylation events in HSF1 activation is unclear. Both S230 and S326 sit in the regulatory domain of HSF1, which has previously been shown to be a region for many post-translational modifications (8, 73). Additionally, it is apparent that the regulatory domain, and S326 specifically, do not regulate nuclear localization or DNA binding of HSF1 (73, 74). Since this domain also does not participate in trimerization, it was tested whether S230 and S326 regulate gene transactivation by interaction with transcriptional machinery. Similar to T527, phospho-null mutants of S230 and S326 had decreased transcriptional activity in the presence of CDK9 (**Fig 8A**). However, only the S326 phospho-null, but not S230A, reduced HSF1 activity in the presence of TFIIB (**Fig 8B**). Similar to T527, neither site had any effect on HSF1 activity in the presence of TBP (**Fig 8C**). It was also observed that the S326A mutant had decreased protein association with CDK9 and TFIIB (**Fig 8D-E**) and the S230A mutant had decreased interaction with CDK9 (**Fig 8F**). Taken together, these results indicate that AKT1-phosphorylated sites functionally contribute to the activation of HSF1 by promoting trimerization and gene transactivation. Considering these sites were exclusive to AKT1 compared to the other known activating kinases, phosphorylation at these sites and the promotion of these activating events likely contributes to the increased potency of HSF1 activation by AKT1.

**Figure 8.**
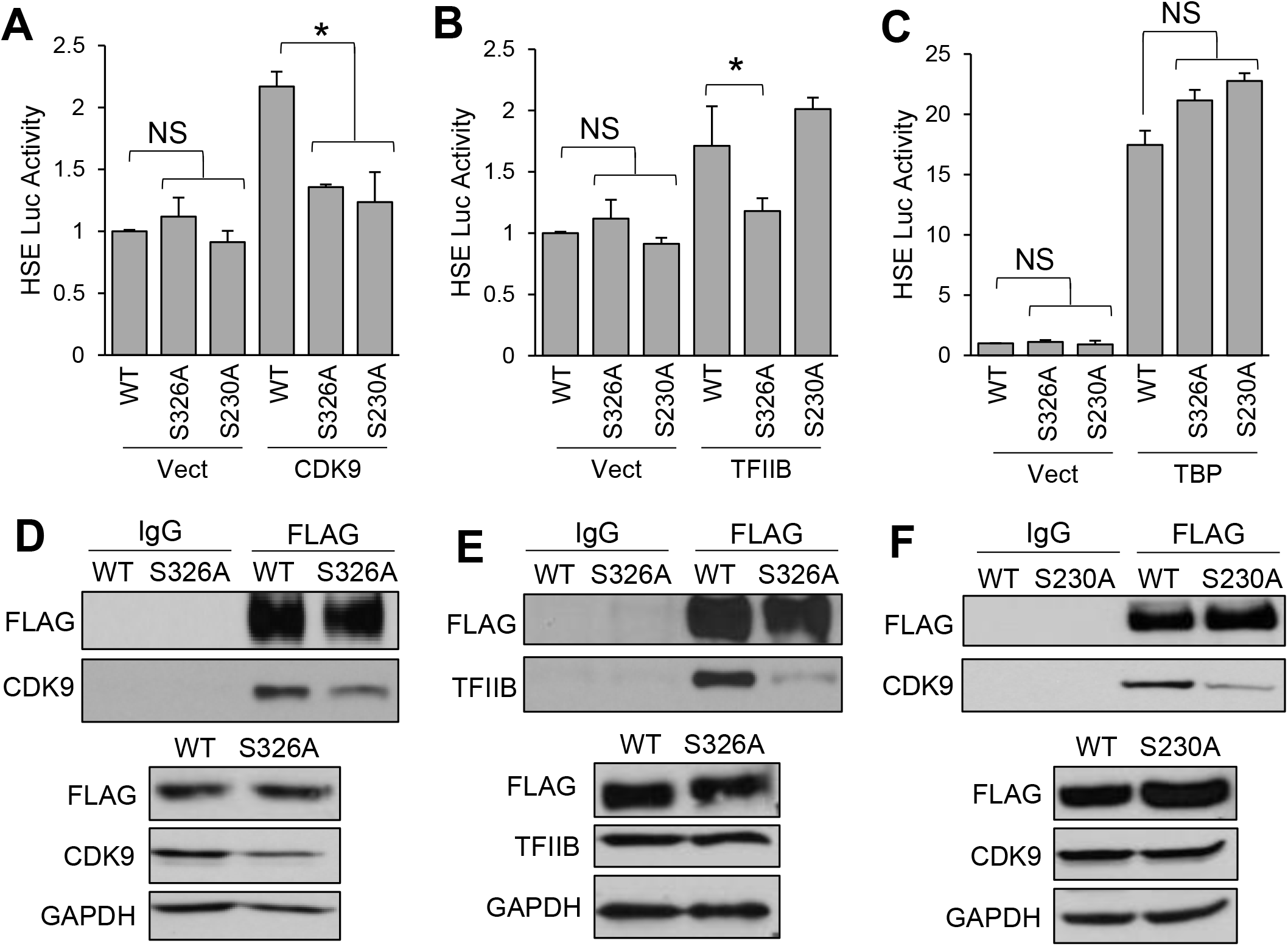
Phosphorylation at S230 and S326 enhances HSF1 association with general transcription factors. A-C) HEK-293 cells were transfected with WT-HSF1, S326A-HSF1, or S230A-HSF1 in the presence or absence of CDK9 (A), TFIIB (B), or TBP (C) along with the HSE luciferase reporter for 48 hrs. Firefly/Renilla ratios were compared across groups. D-F) HEK-293 cells were transfected with FLAG-tagged WT-HSF1, S326A-HSF1, or S230A-HSF1 for 48 hrs. Cell lysates were subjected to immunoprecipitation with control IgG or FLAG antibodies and subjected to immunoblotting with indicated antibodies. A-C were completed in triplicate and analyzed using one-way ANOVA and Tukey’s post-hoc test. Data is presented as mean ± SEM. *P<0.05.

## Discussion

The heat shock response is primarily regulated by increasing the transcriptional activity of HSF1 in response to heat stress. In order for HSF1 to become transcriptionally active, it must overcome cytoplasmic sequestration followed by nuclear localization, trimerization, DNA binding, phosphorylation, and gene transactivation. The regulation of HSF1 activity by phosphorylation has been debated but what is generally accepted is that S326 phosphorylation is important for HSF1 activity, a site that was initially identified from an HSF1 phospho-null screen (8). Considering the importance S326 phosphorylation, a better understanding of the mechanisms regulating phosphorylation at this residue would likely provide insight on how to manipulate HSF1 activity in disease conditions wherein HSF1 is dysregulated. Phosphorylation at S326 has been shown to be mediated by several kinases including mTORC1 (33, 37), AKT1 (15), MEK1 (38), and p38 (34). The current study additionally indicates that AKT2 has the capability to phosphorylate S326 (**Fig 1**). The involvement of these kinases appears to be context dependent. The current study indicated that the context-dependent nature of these kinases may stem from the unique pattern of residues phosphorylated by each kinase. This study indicates mTORC1, MEK1, and p38 induce HSF1 activity to a similar level but phosphorylate HSF1 primarily at S326 as well as S363, a known inhibitory site (75, 76), S221, also thought to be an inhibitory site (74), or at S241 and S344, which are two novel phosphorylation sites with unknown function. However, AKT1 appeared to be a more potent inducer of HSF1 activity that likely stems from its ability to phosphorylate an additional three residues identified here that all functionally promote HSF1 activation. The PI3K pathway has been linked to HSF1 activity as far back as 2000 (59, 77) but whether this was an indirect regulation or a direct regulation was unclear until our previous study indicating AKT1 directly activates HSF1 (15). The current study indicates that AKT1 also participates in heat stress-induced activation of HSF1 (**Fig 2**) and that AKT1-mediated activation is independent of mTORC1 (**Fig 1**).The current study extends our previous findings to identify precisely how AKT1 regulates HSF1 activity through phosphorylation. Additionally, this study indicates for the first time the functional role in HSF1-induced transcription for each of these phosphorylation events by enhancing trimerization and gene transactivation.

Upon heat stress, HSF1 can be observed to undergo hyperphosphorylation. Hyperphosphorylation was initially thought to be a part of the HSF1 activation process (10, 12, 78)). A more recent study indicated that an HSF1 mutant wherein all of the serine residues within the regulatory domain are phospho-null has greater activity compared to WT-HSF1 and this mutant did not undergo classic hyperphosphorylation (74), suggesting that hyperphosphorylation is actually suppressive to HSF1 activity. There were also early indications that extensive phosphorylation can suppress HSF1 activity (79). However, results from the current study indicate that phosphorylation on specific residues enhance HSF1 transcriptional activity by mediating specific steps in the process of activation. Thus, it seems that the model for HSF1 activation involves phosphorylation at specific residues but then hyperphosphorylation leads to inactivation of HSF1. This model would presume that activation phosphorylation events occur prior to hyperphosphorylation, although there is currently no published evidence for this temporal relationship. This model is also supported by the numerous studies indicating that specific phosphorylation on S326 is important for HSF1 activity. While this model is intriguing as a potential negative feedback on HSF1, the mechanism for hyperphosphorylation, and associated kinases involved, remain to be uncovered.

Phosphorylation at S326 has been used as a biomarker for transcriptionally active HSF1 in many studies since the identification of this post-translational modification. The current study confirms previous studies that multiple kinases can phosphorylate this residue, which was associated with increased HSF1 activity (**Fig 4**). However, the current study also identified that AKT1 was a stronger inducer of HSF1 activity than the other kinases tested despite S326 being phosphorylated by all kinases tested. This apparent paradox prompts the question of whether S326 phosphorylation is a reliable marker for HSF1 activity. Other reports indicate several inhibitory phosphorylation sites on HSF1, most notably S303/307 and S363 (75, 76, 80–84). Several of these studies provide evidence that phospho-null mutations at these inhibitory sites can increase HSF1 activity but S326 can presumably still be phosphorylated under these circumstances. The current study indicates S326 phosphorylation actively participates in HSF1-induced gene transactivation giving support that it can serve as a marker for activity. Other studies have also observed S326 phosphorylation can still be used as a marker in disease contexts, such as S326 phosphorylation serving as a biomarker for ovarian cancer (39). Thus, based on the current and previous studies, it is likely that S326 phosphorylation is a marker for transcriptionally competent HSF1 but may not fully indicate the intensity of activity.

The potency for AKT1-mediated activation of HSF1 observed in the current study advances our understanding of this relationship in disease settings wherein HSF1 is dysregulated. In particular, HSF1 is well known to be hyperactivated in several cancer types (14, 15, 17–20, 22, 36, 38, 39, 46). Hyperactivation of HSF1 in cancers likely stems from the presence of several cellular stressors in cancer cells and the tumor microenvironment. The kinases known to activate HSF1 and tested in this current study all have been shown to be dysregulated in several cancers. While AKT1, mTORC1, MEK1, and p38 can all activate HSF1, the current study indicates that activity of only AKT1 and mTORC1 maintain a strong association with HSF1 activity in tumors (**Fig 4**). It is likely that AKT1 activity maintains a strong association with HSF1 activity due to the mechanisms uncovered in this study that AKT1 can phosphorylate multiple residues that promote HSF1 activity (**Fig 5–8**). While it is possible that AKT1 can increase mTORC1 activity, this study also indicates that AKT1-driven activation of HSF1 is independent of mTORC1 (**Fig 1**). The more likely explanation for the association between HSF1 activity and mTORC1 activity in these cancer datasets is that HSF1 has been shown to enhance mTORC1 activity via suppression of JNK (37). This would likely explain the stronger association between HSF1 and mTORC1 if HSF1 is acting upstream of mTORC1.

These studies together indicate that HSF1 activity is regulated by phosphorylation at specific residues. In particular, phosphorylation at the canonical activate site of S326 promotes gene transactivation by enhancing the association between HSF1 and CDK9/TFIIB. Furthermore, AKT1 can phosphorylate additional residues that further enhance HSF1 activation including T142, S230, and T527. Phosphorylation at T142 has been previously observed by CK2 (64) and we show here that T142 phosphorylation enhances trimerization of HSF1 in response to AKT1 or CK2. Phosphorylation at S230 has been observed by CaMKII (65) and we show here that S230 phosphorylation by AKT1 enhances HSF1 association with CDK9 to promote transcription. Phosphorylation at T527 has never been reported and we observed AKT1-mediated phosphorylation of T527 enhances the association between HSF1 and CDK9/TFIIB to promote transcription. These studies illuminate further the complex regulation of HSF1 activation and, in concert with previous reports, likely indicate that HSF1 is activated by phosphorylation at a few specific residues, such as those reported here, but that hyperphosphorylation is likely suppressive. It will be important to identify the proteins involved in hyperphosphorylation-mediated inactivation of HSF1 as these could be therapeutic targets in HSF1-dysregulated diseases. Additionally, a greater understanding of HSF1 post-translational modifications in response to various cellular stressors would likely indicate key events required for consistent HSF1 activation.

## Acknowledgements

This work was supported by funds from the National Cancer Institute (K22CA207575, RLC). We would like to acknowledge the Laboratory for Biological Mass Spectrometry at Indiana University for help with performing the mass spectrometry experiments and the associated data analysis.

## Nonstandard Abbreviations

GSEA: Gene Set Enrichment Analysiss
HSE: Heat shock element
HSF1: Heat shock factor 1
TCGA: The Cancer Genome Atlas

## Author Contributions

RLC conceived the idea and supervised the project. WCL collected and analyzed data for trimerization, ChIP, co-immunoprecipitation, and some reporter assay experiments as well as generated all mutant constructs used in this study. RO and HR performed all kinase assays as well as some immunoblotting and reporter assay experiments. CJ and NH performed some immunoblotting, reporter assays, and qPCR experiments. MB performed size-exclusion chromatography. All authors assisted with data discussion and manuscript preparation. WCL and RLC wrote the manuscript with edits provided by all authors.

