## Supplemental Information for "AKT1 mediates multiple phosphorylation events that functionally promote HSF1 activation"

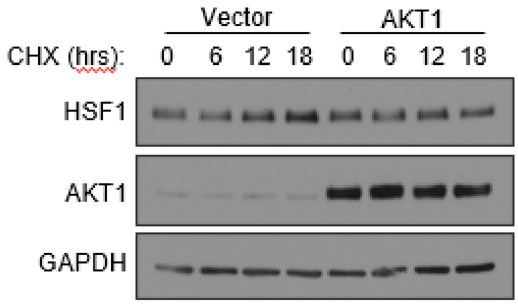


**Supplemental Figure 1: HSF1 protein stability is not affected by AKT1 presence.** HEK-293 cells were transfected with either an empty vector or AKT1 followed by treatment with cycloheximide (10 ug/mL) for up to 18 hrs. Cells were then lysed and total protein subjected to immunoblotting with indicated antibodies. HSF1 protein stability was not altered in the presence of AKT1.

**Supplemental Table 1: AKT1 has a unique phosphorylation pattern on the HSF1 protein**

| Kinase | HSF1 Peptide | Peptide Start | P-site |
|---|---|---|---|
| AKT1 | SVTKLL**pT**DVQLM | 136 | T142 |
| | **pS**LEHVHGSGPY | 230 | S230 |
| | TGSEPPKAKDP**pT**VS | 516 | T527 |
| AKT2 | **pS**APSPAYSSSSLY | 241 | S241 |
| | TDARGHTDTEGRPP**pS**PPPTSTPEKCL | 349 | S363 |
| P38 | **pS**APSPAYSSSSLY | 241 | S241 |
| | TDARGHTDTEGRPP**pS**PPPTSTPEKCL | 349 | S363 |
| mTORC1 | MLNDSGSAH**pS**MPKYSRQF | 212 | S221 |
| | **pS**APSPAYSSSSLY | 241 | S241 |
| | IDSILRESEPAPA**pS**VTAL | 331 | S344 |
| MEK1 | TGSEPPKAKDP**pT**VS | 516 | T527 |

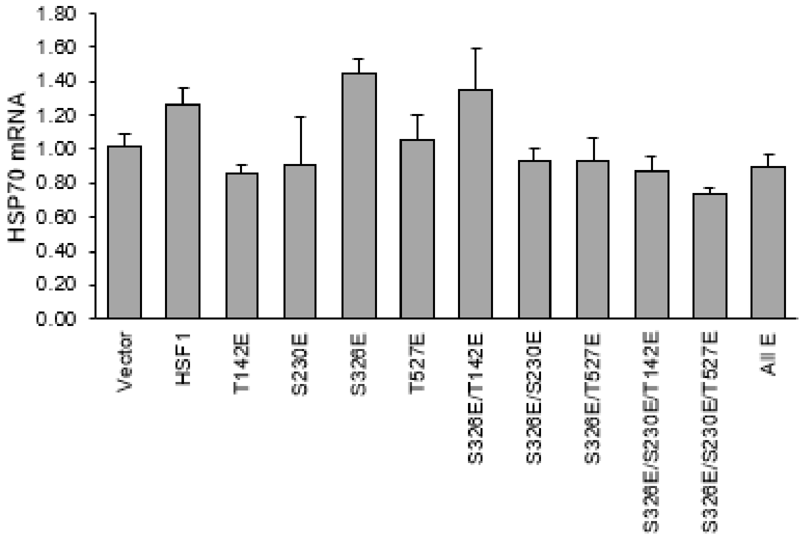


**Supplemental Figure 2: Serine to Glutamate Mutation does not result in a Phospho-mimic for HSF1 phosphorylation sites.** Wild-type HSF1 underwent mutagenesis at the indicated residues. An empty vector, wild-type HSF1, and each of these mutants were expressed in HEK-293 cells. Total RNA was subjected to RT-qPCR for HSP70, an HSF1 target gene. None of these phospho-mimic constructs were able to increase HSF1 activity over the wild-type suggesting they do not sufficiently mimic these residues being phosphorylated.

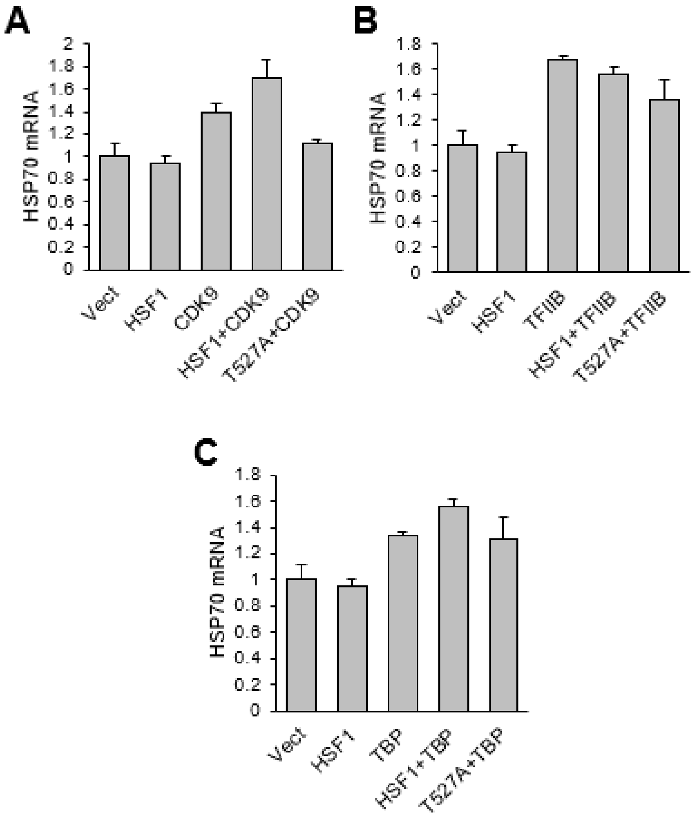


**Supplemental Figure 3: HSF1 activity in the presence of CDK9 and TFIIB is reduced with T527A mutation.** Wild-type HSF1 was mutated to give T527A mutation. Empty vector, wild-type HSF1, or T527A were expressed with or without CDK9 (A), TFIIB (B), or TBP (C). Total RNA was subjected to RT-qPCR for HSP70, a direct target gene of HSF1. T527A mutation significantly reduced HSF1 activity in the presence of CDK9 and TFIIB.
